# Chemogenetic stimulation of a retinal circuit activates brain noradrenergic neurons, prevents apoptosis and suppresses depression-like behaviors

**DOI:** 10.1101/2021.04.20.440684

**Authors:** Hannah E Bowrey, Morgan H James, Mohsen Omrani, Aida Mohammadkhani, G Aston-Jones

## Abstract

A chronically dysregulated locus coeruleus (LC) system gives rise to mood disorders. In particular, prolonged LC hyperactivity often underlies stress disorders, whereas chronically reduced function often underlies symptoms of depression. Owing to its location deep in the brainstem, LC is difficult to access which limits translational approaches that involve manipulating these neurons. Here, we circumvent this problem by utilizing the retina as a chemogenetic target for commandeering a multsynaptic circuit to LC. We show that activation of this pathway can prevent depression-like behavior and associated pathology of the LC-noradrenergic (NA) system caused by light deprivation. Additionally, we show that melanopsin-containing intrinsically photosensitive retinal ganglion cells (ipRGCs) are the likely cell type responsible for initiating activity in this pathway. By capitalizing on this ocular route of designer receptor delivery, we demonstrate a novel, minimally-invasive method for manipulating this deep-brain circuit that is relevant to a wide array of neuropsychiatric disorders.

---

Previous findings have critically implicated the noradrenergic (NA) brain nucleus locus coeruleus (LC) in numerous neuropsychiatric processes and diseases^1–3^, including depression^4^. Thus, the selective manipulation of LC represents an important translational target for treating psychiatric disease. However, due to LC’s deep location and small size, suitable approaches for its manipulation either by genetic vector-based control or by electrode-based stimulation are lacking.

Designer receptor technologies, such as DREADDs (Designer Receptors Exclusively Activated by Designer Drugs) hold substantial translational potential. However, current technology requires the direct injection of a DREADD vector directly into the brain, limiting its clinical utility; this is particularly true for disorders that involve deep brain structures such as LC. The retina presents a unique avenue for DREADD transduction as is can be accessed from the periphery. Additionally, adeno-associated virus (AAV), a vector utilized to deliver the DREADD gene, has an excellent safety profile and provides long-term expression in the human retina^5^ so that DREADDs can be easily transduced into retinal ganglion cells (RGCs) via a simple intraocular injection^6^. This technique is rapid in the laboratory, and cost-effective and safe in the clinic, thus representing an excellent minimally invasive method for delivering DREADDs to the central nervous system.

We previously described a circuit from suprachiasmatic nucleus (SCN) to locus coeruleus (LC) that provides circadian regulation of arousal; this circuit utilizes dorsomedial hypothalamus (DMH) as a relay^7^. Because the retina provides photic input to SCN, we propose a multisynaptic circuit (retina-SCN-DMH-LC) for mediating at least some effects of light on mental functions such as mood^4^; also, alterations in this circuit may contribute to pathological changes observed in LC following aberrant lighting conditions. For example, light deprivation induces a depressive-like phenotype, apoptosis in LC-NA neurons, and loss of NA cortical fiber density in frontal cortex^8^. Due to its indirect input from retina, we reasoned that DREADD- activation of retina may stimulate LC and prevent light deprivation-induced LC apoptosis, cortical NA fiber loss, and depression-like behavior.

We analyzed evidence to determine whether this pathway, which we have termed the Photic Regulation of Arousal and Mood (PRAM) pathway (Fig 1.A.), is functional. First, we expressed either neuron-specific (hSyn) promoter-driven excitatory DREADDs (AAV2-hsyn-hM3Dq; hereafter referred to as hM3Dq) or control virus (AAV2-hsyn-GFP; hereafter referred to as GFP) in retinae of Sprague Dawley rats via intravitreal injection (IVI) (Fig 1.B.). Fos activation of DREADD-labeled cells following CNO administration indicated retinal activation in hM3Dq animals (Fig 1.C.). DREADD-labelling was restricted to the ganglion cell layer (GCL) of the retina, indicating that RGCs were preferentially transduced (Fig. 1.D). DREADD activation (by CNO) increased the sensitive state of the retina as measured by ERG (Fig. 1 E,F.). In addition, DREADD-activation of retina induced Fos expression in retina, SCN, DMH and LC of hM3Dq rats compared with control littermates (Figs 1.G,H,I,J), with at least a doubling of active cells in all PRAM structures (all *p*’s <0.0001). Impulse activity of LC neurons *in vivo* (isoflurane anesthesia) was recorded before and after CNO-mediated retinal DREADD activation, while eyes were protected from light. DREADD-mediated activation of the retina significantly increased the firing rate of LC cells above baseline activity (from 1.91 ± 0.28 Hz to 3.19 ± 0.52 Hz, *p* = 0.01); no such changes were observed in littermate GFP controls (Figs 1. K,L.).

**Figure 1.**
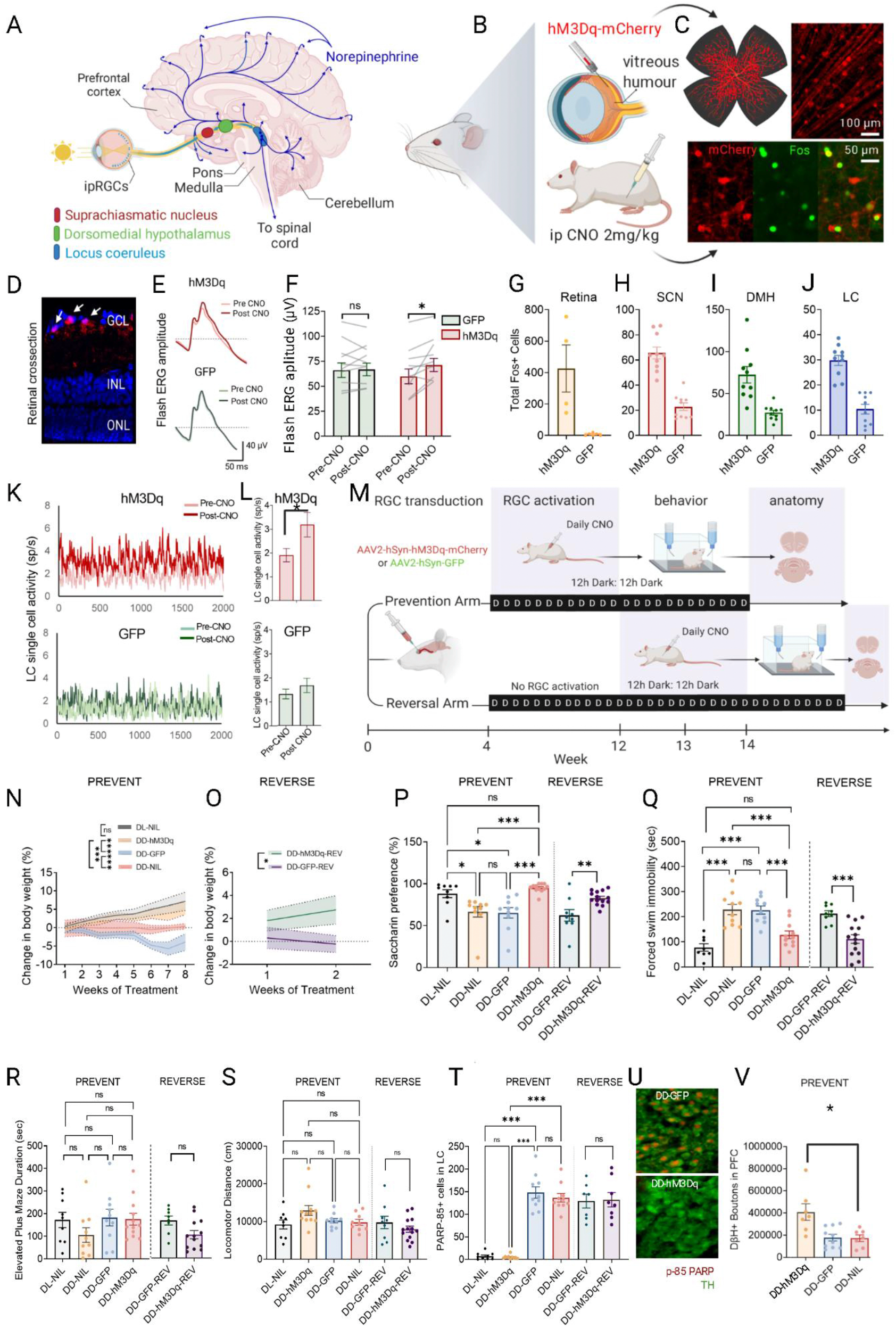
Chemogenetic retinal activation prevents and reverses light deprivation-induced depression-like behaviors and LC-NA pathology. (A) Schematic of the photic regulation of arousal and mood (PRAM) pathway. (B) Injection strategy for delivery and activation of DREADDs. (C) Expression of DREADDs and colocalized retinal Fos following CNO administration. (D) Z-Stack of retina cross-section showing DREADD expression (red) in ganglion cell layer (GCL). (E) Full flash electroretinogram (ERG) typical trace of individual hM3Dq (upper) and GFP (lower) animals, pre- and post-CNO administration. (F) ERG amplitutdes; *p < 0.05. (G – J) Number of Fos-active cells in key PRAM structures (G:retina; H:suprachiasmatic nucleus (SCN);I: dorsomedial hypothalamus (DMH); and J:locus coeruleus (LC)) following retinal DREADD activation (2mg/kg CNO, ip). (K) Single unit recording of a sample LC neuron before and after CNO administration (2 mg/kg ip) in a rat expressing hM3Dq in retinae. (L) CNO did not significantly change LC activity in rats expressing control virus (GFP) in retinae (p=0.49; Wilcoxon ranksum test), but tonic LC activity was increased following CNO in animals expressing retinal hM3Dq (*p* = 0.01, Wilcoxon ranksum test). (M) Schematic of behavioral experiment showing prevention (PREV: 8wks of daily CNO concurrent with constant darkness (DD)) and reversal paradigms (REV: 2wks of daily CNO concurrent with DD, immediately following 8 weeks of DD without CNO). (N) Retinal activation prevents DD-induced relative weight loss. (O) Retinal activation reverses DD-induced relative weight loss. (P) Anhedonia-like behavior (as seen by decreased saccharine preference) is prevented and reversed by retinal activation. (Q) Despair-like behavior (as measured by forced swim test) is prevented and reversed by retinal activation. (R) Retinal activation has no significant effect on anxiety-like behavior in the elevated plus maze. (S) There was no effect of retinal activation on locomotor behavior in an open field, indicating our results cannot be explained simply by increased locomotor activity. (T) Retinal activation prevented, but did not reverse, apoptosis in LC cells (P-85 PARP). (U) Apoptosis (red nuclei, reflecting P-85 PARP) in DDhM3Dq LC NA neurons (upper) vs apoptosis in DD- GFP LC. LC NA neurons stained green with an antibody to tyrosine hydroxylase. (V) Retinal stimulation increased DβH+ boutons in prefrontal cortex (PFC) in the prevention arm. Error bars are s.e.m. *p < 0.05, **p < 0.01, ***p < 0.001

We then asked whether chemogenetic activation of the PRAM pathway would prevent and/or reverse the development of depression-like behavior induced by light deprivation (constant dark; DD). We employed classic rodent tests to estimate depression-like behavior, including weight loss, the saccharin preference test (SPT), and the forced swim test (FST)^9^. We expressed either hM3Dq (hereafter referred to as DD- hM3Dq, *n* = 12), control virus (DD-GFP, *n* = 10), or no virus (DD-NIL, *n* = 10) in both retinae of rats maintained in DD for 8 weeks. They were then assigned to either the *prevention arm* (PREV) or the *reversal arm* of the study (REV), the latter better modeling potential clinical chronology (Fig. 1M; Table 1.). All animals that received either hM3Dq or GFP vectors intraocularly, also received daily CNO (2 mg/kg, i.p.) at the beginning of their active period (19:00h each evening). PREV animals (both hm3Dq and GFP) received CNO concurrently with 8 weeks of DD. REV (both hM3Dq and GFP) animals received 2 weeks of daily CNO (during DD) following 8 weeks of DD. An additional PREV control group was maintained in chronic DD but received no ocular vector injection or CNO (DD-NIL, *n* = 10), and a final PREV group was maintained in standard 12h dark: 12h light (DL) conditions with no ocular or i.p injections (DL-NIL, *n* = 9). In the PREV and REV arms, daily DREADD-mediated activation of retinae prevented or reversed, respectively, the development of multiple light deprivation-induced depression-like behaviors relative to control littermates. Specifically, daily DD-hM3Dq (PREV and REV) activation prevented or reversed the relative weight loss observed in DD-NIL and DD-GFP PREV and REV groups, respectively (Fig.1.N,O.). Additionally, daily DREADD-activation of retinal cells decreased anhedonic-like behavior, as measured by a saccharin preference test (SPT), and reduced despair-like behavior as measured by a forced swim test (FST), relative to littermate controls (Fig.1.P,Q). DD-hM3Dq (PREV and REV) activation had no significant effect on elevated plus maze activity (EPM) or locomotor activity, and DD-hM3Dq (PREV) was similar to the DL-NIL PREV group in both measures. (Fig.1.R,S). Together, these data indicate that activating the PRAM pathway can both prevent and reverse the development of depressive-like behaviors caused by chronic light deprivation, without affecting anxiety-like or locomotor behaviors.

**Table 1.**
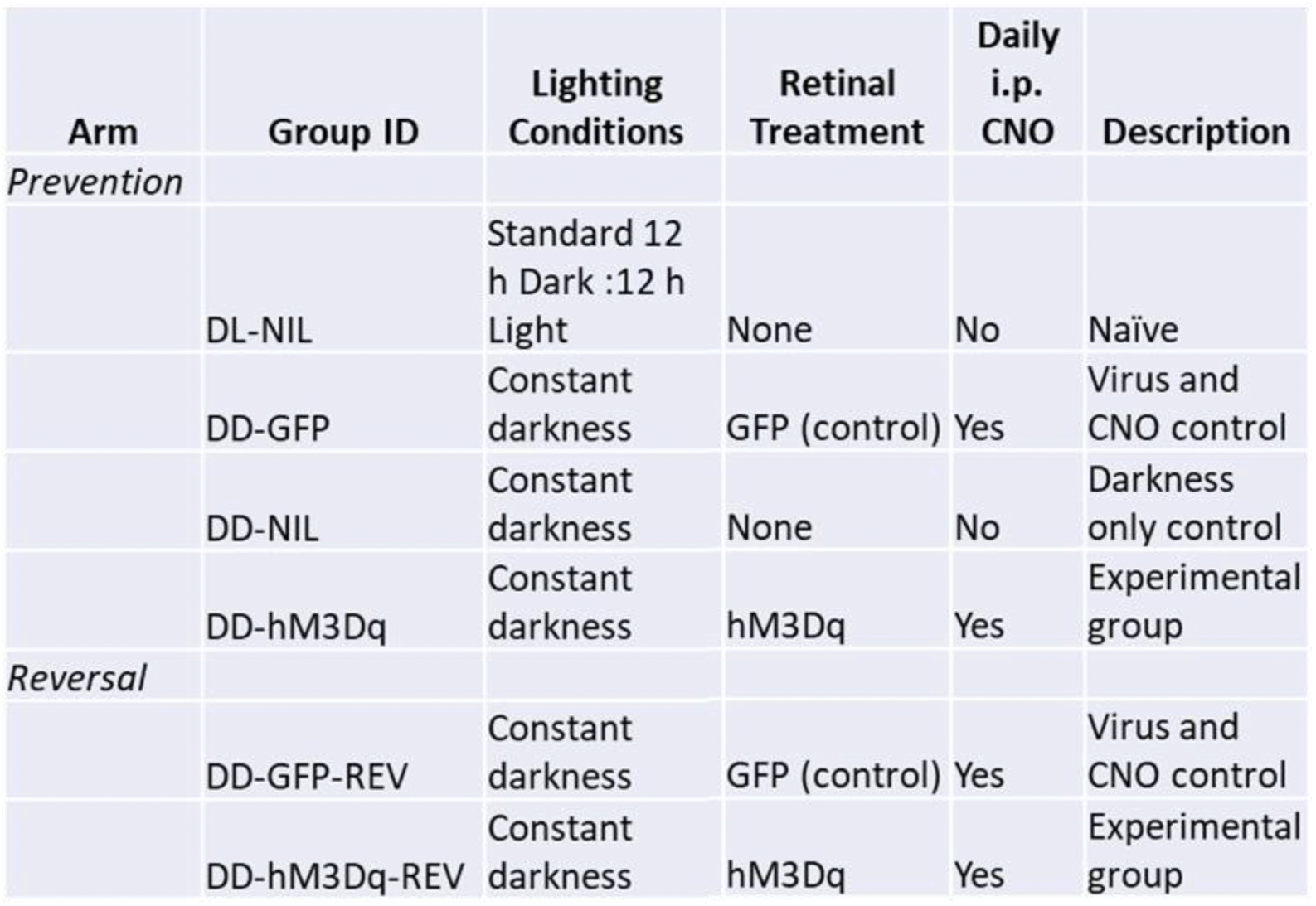
Arm allocations, group ID, lighting conditions, and treatment of animals in Experiment 1.

| Arm | Group ID | Lighting Conditions | Retinal Treatment | Daily i.p. CNO | Description |
| --- | --- | --- | --- | --- | --- |
| *Prevention* |  |  |  |  |  |
|  | DL-NIL | Standard 12 h Dark :12 h Light | None | No | Naïve |
|  | DD-GFP | Constant darkness | GFP (control) | Yes | Virus and CNO control |
|  | DD-NIL | Constant darkness | None | No | Darkness only control |
|  | DD-hM3Dq | Constant darkness | hM3Dq | Yes | Experimental group |
| *Reversal* |  |  |  |  |  |
|  | DD-GFP-REV | Constant darkness | GFP (control) | Yes | Virus and CNO control |
|  | DD-hM3Dq-REV | Constant darkness | hM3Dq | Yes | Experimental group |

We also asked whether daily retinal DREADD activation would prevent and/or reverse the apoptosis in LC-NA neurons that normally occurs with chronic light deprivation (8). Daily DD-hM3Dq activation prevented but did not reverse apoptosis in LC (as measured by p-85 fragment of Poly (ADP-ribose) polymerase; PARP), which was abundant in littermate DD controls (Fig 1. T,U).). In agreement with these findings, we observed a significant reduction in dopamine-β-hydroxylase-positive (DβH+) fibers in prefrontal cortex (PFC) of DD controls, compared with DD-hM3Dq and DL-NIL, in the PREV arm (Fig. V.). Thus, chronic light deprivation leads to apoptosis in LC and a reduction in LC-projecting DβH+ PFC fibers, which is prevented, but not reversed, by daily hM3Dq activation of retinae.

We hypothesized that LC function and NA cortical innervation play a critical role in the rescue of normal affective behavior by retinal stimulation in light deprived animals. We either ablated central DβH+ projections using N-(2-chloroethyl)-N-ethyl-2-bromobenzylamine (DSP-4; hereafter referred to as ABLATE group) or selectively inhibited LC-NA cells via direct insertion of an AAV-DIO-hM4Di-EGFP vector into LC of TH::Cre rats (hereafter referred to as INHIBIT group) during DD and retinal stimulation (Fig.2A.). ABLATE animals were injected with DSP-4 (50 mg/kg ip) plus either hM3Dq or GFP IVI (DD-hM3Dq-DSP4, *n* = 9; and DD-GFP-DSP4, *n* = 9). An additional group received vehicle for DSP-4 (ip) plus intraocular hM3Dq (DD- hM3Dq-VEH, *n* = 11). INHIBIT animals were TH::Cre rats that were intraocularly injected with hM3Dq, plus DIO-hM4Di locally microinjected into LC (DD-hM3Dq-hM4Di; *n* = 9). Controls for INHIBIT were WT with DIO-hM4Di or control (GFP) virus directed into LC, respectively (DD-hM3Dq-CTRL; *n* = 5).

The blockade of anhedonic-like behavior (SPT) normally seen after Gq PRAM activation (described above) was prevented in both ABLATE and INHIBIT groups, compared with litter-mate controls (Fig. 2D.). In addition, blockade of despair-like behavior (FST) normally seen after Gq PRAM activation (described above) was completely prevented if LC projections were ablated or if LC-NA cells were inhibited (Fig. 2E). Neither ablation of central NA projections by DSP4 (Fig. 2B) nor inhibition of LC-NA activity by hM4Di (Fig. 2C.) prevented the normal weight-gain in animals that received hM3Dq-driven retinal stimulation during DD, likely indicating that LC does not play a necessary role in this phenomenon. Neither LC ablation nor inhibition had a significant effect on EPM (anxiety; Fig.2F.) or locomotor activity (Fig.2 G.) in animals that also received retinal PRAM activation. Taken together, these data indicate that LC is necessary for prevention of many, but not all, of the depression-like behaviors by retinal activation.

**Figure 2.**
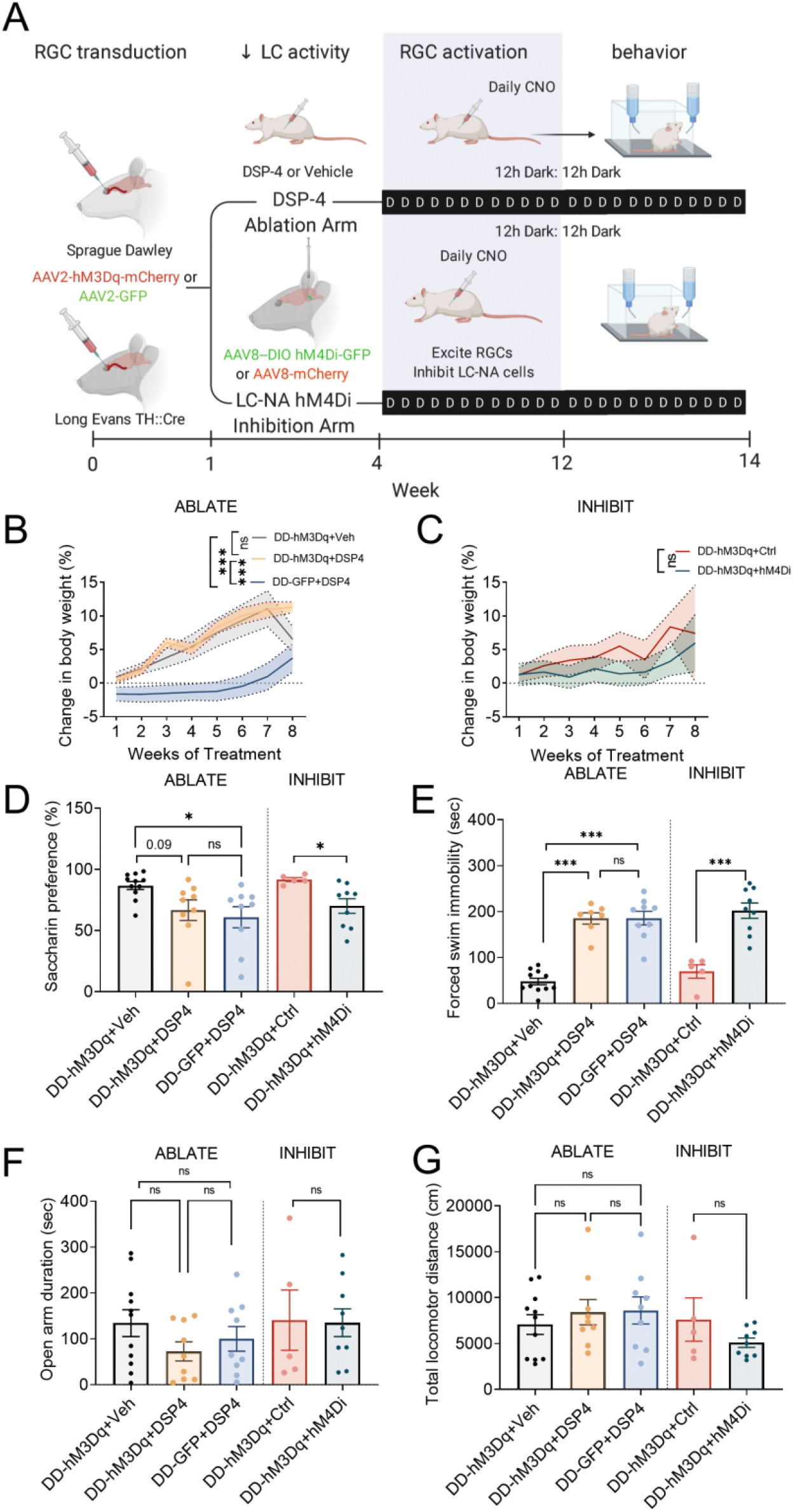
Locus coeruleus activity is required for retinal stimulation to prevent depression-like behavior. (A) Schematic illustrating ablation of DβH+ fibers (using DSP-4) or inhibition of LC (using hM4Di DREADD) to test the necessity of LC for efficacy of PRAM stimulation (intraocular hM3Dq) to prevent depression-like behaviors. (B) Ablation of DβH+ fibers does not prevent the normalization of weight gain by PRAM stimulation. (C) Inhibition of LC signaling does not prevent the normalization of weight gain by PRAM stimulation. (D) LC-NA projections and activity are required for prevention of anhedonia-like response to DD (SPT). (E) LC-NA projections and activity are required for prevention of the despair-like endophenotype in response to DD. (F & G) There were no significant effects of DSP4 lesion or LC chemogenetic inhibition on anxiety (EPM) or (G) locomotor activity (open field). Error bars are s.e.m. *p < 0.05, **p < 0.01, ***p < 0.001

We investigated the role of intrinsically photosensitive RGCs (ipRGCs) ^10,11^ in the above sets of results. These melanopsin-containing ipRGCs project directly and strongly to SCN ^12^, and have been critically implicated in light-mediated mood modulation ^13,14^. They are primarily involved in non-image-forming functions, such as circadian entrainment ^15^ and pupil dilatory response ^16,17^, and thus offer a tantalizing clinical route for manipulation of the PRAM pathway while minimizing effects on image-forming vision ^4^. We asked whether a reduction of ipRGC activity may underlie depressive-like behavior and LC pathology in DD rats.

We designed a saporin toxin conjugated to a melanopsin antibody to selectively ablate ipRGCs (similar to ^18,19^), which was injected bilaterally (IVI) in each animal, ablating ~50% of ipRGCs (Fig. S1). Control animals were similarly injected with 0.01M PBS. Animals from both experimental and control groups were maintained in standard DL conditions for 8 weeks following these IVI injections (DL-MEL-SAP, *n* = 9; DL- VEH, *n* = 9, respectively). For comparison, a third group was bilaterally injected IVI with 0.01M PBS, but was placed in DD (DD-VEH, *n* = 10; Fig. 3A.).

****Figure 3.**
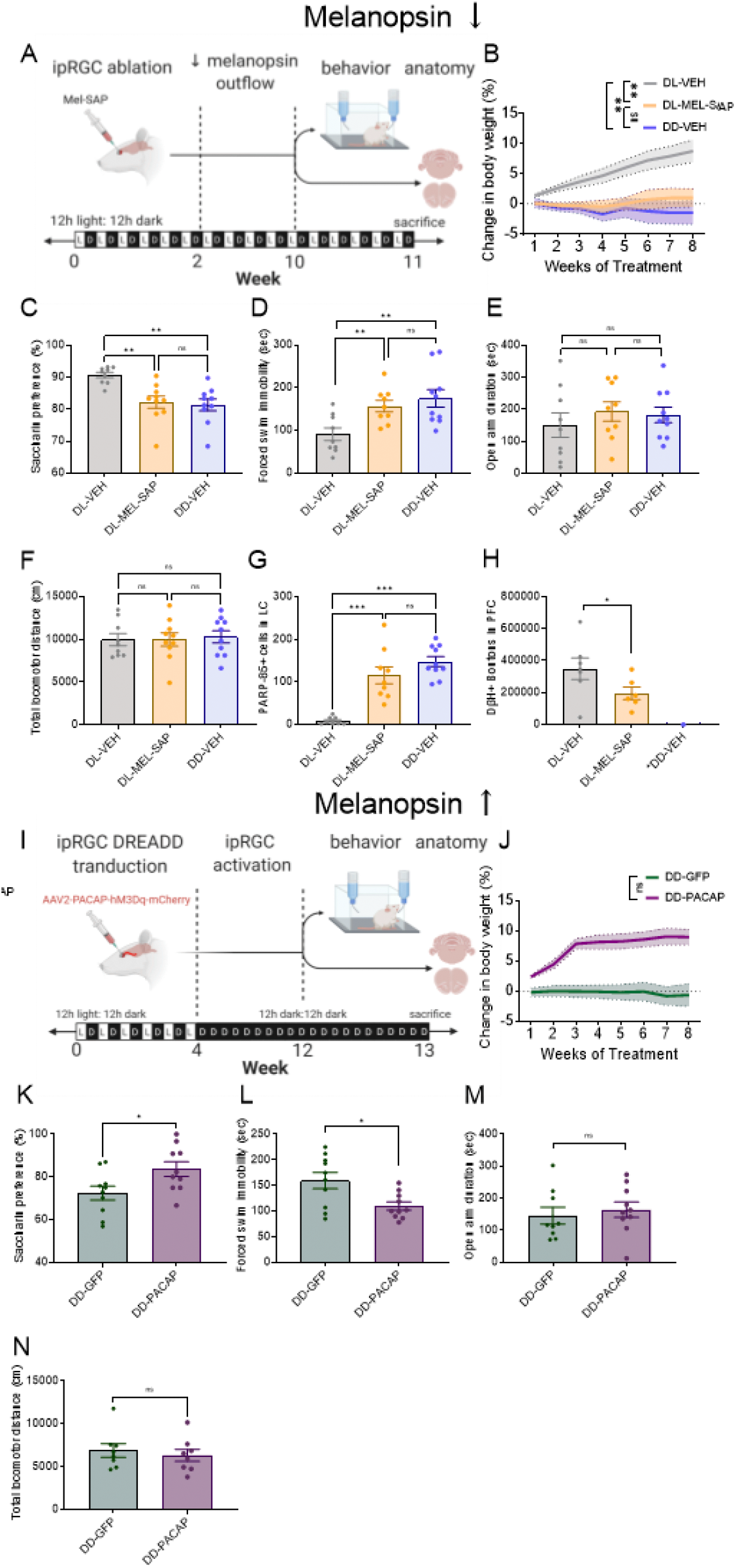
Effects of ipRGC ablation and stimulation on the development of depression-like behavior. (A) Schematic demonstrating paradigm using ablation of ipRGCs. (B) ipRGC ablation prevents normal weight gain. (C) Saccharin preference is reduced by ablation of ipRGCs. (D) Forced swim is decreased. by ablation of ipRGCs. (E) ipRGC ablation had no significant effect on EPM open arm time. (F) There was no effect on locomotor behavior after ablation of ipRGCs. (G) ipRGC ablation induced apoptosis in LC cells (p85 PARP). (H) ipRGC ablation reduced the number of DβH+ boutons in prefrontal cortex (PFC) (DD-VEH counts are ongoing). (I) Schematic demonstrating method for activating ipRGCs using PACAP-promoter hM3Dq DREADD. (J) ipRGC activation by PACAP hM3Dq facilitates normal weight gain. (K) Decreased saccharin preference in DD is prevented by activating ipRGCs. (L) Decreased swim in DD is prevented by activating ipRGCs. (M) ipRGC activation has no significant effect on EPM open arm time. (N) ipRGC activation has no significant effect on locomotor behavior. Error bars are s.e.m. *p < 0.05, **p < 0.01, ***p < 0.001

Specific ablation of ipRGCs (DL-MEL-SAP) induced significant weight loss relative to the DL-VEH controls; this weight loss was equivalent to that exhibited by control rats kept in constant darkness (DD-VEH) (Fig 3B). Anhedonia-like behavior (SPT) was greater in DL-MEL-SAP compared with DL-VEH (Fig. 3C.) and was comparable to DD-VEH animals. Ablation of ipRGCs also caused despair-like behavior as measured by FST (Fig. 3D), whereby time spent immobile was greater in DL-MEL-SAP animals compared with DL-VEH animals. This again was comparable to DD-VEH animals. We found no effect of ipRGC ablation on EPM (anxiety) or locomotor activity (Fig. 3E, F.). Taken together these data indicate that chronically reducing the outflow of ipRGCs has a pro-depressive effect similar to chronic DD, which is not due to anxiety or general locomotor deficits. DL-MEL-SAP was also associated with increased apoptosis in LC-NA neurons (PARP). This was indistinguishable from PARP-apoptosis in DD-VEH, and in contrast to DL-VEH that showed no evidence of LC-NA apoptosis (Fig. 3G). In line with the robust apoptotic response of LC, we observed a reduction in DβH+ fibers in PFC in DL-MEL-SAP relative to DL-VEH subjects (Fig. 3H.).

Melanopsin cells co-express the peptide pituitary adenylate-cyclase-activating polypeptide (PACAP) ^20,21^. To selectively enhance ipRGC activity during DD, we developed a PACAP-promoter driven DREADD (AAV- 2-PACAP-hM3Dq-mCherry; DD-PACAP). Animals were binocularly injected with this vector (*n* = 9), and a negative control group was injected in an identical manner but with a control virus (DD-GFP; *n* = 9) (Fig. 3I.). DD-PACAP activation produced behavior that closely followed the same pattern as DD-hM3Dq (from Fig.1): animals experienced normal weight-gain (Fig. 3 J), greater hedonic-like behavior (Fig 3.K), reduced despair (Fig. L) and no significant differences in behavior on EPM (Fig. 3M.) or locomotor activity (Fig. 3N), compared with litter-mate controls. These data indicate that ipRGCs are likely the cells initiating the PRAM pathway and are influencing the endophenotypic behaviors associated with the prevention of depression-like behavior by retinal stimulation.

## Discussion

Here we show that PRAM stimulation of LC can prevent depression-like behavior in an animal model induced by chronic light deprivation and avoid the need for direct brain injections. Our results confirm a functional multisynaptic circuit from retina to LC. We found that the PRAM pathway is effective for both the prevention as well as the reversal of depressive-like behaviors. This finding is supported by numerous previous studies that demonstrate a role for light exposure on mood. Our findings indicate that positive affective response to light stimulation may be due to activity of LC neurons, which we find essential for preventing and reversing despair- and anhedonia-like behaviors.

Our findings support previous basic ^8,22^ and clinical ^23–26^ studies that report a deficit of NA-LC function in depression-like behavior. We build on these studies by demonstrating that activation of the PRAM pathway (including LC) prevents this deficit in an LC-dependent manner. Our findings indicate that ipRGCs are a likely cell type responsible for this LC activation, as selective ablation of these neurons mimicked effects of light deprivation on LC pathology and associated depressive-like symptoms. This is in line with previous findings that wavelengths of light that activate human ipRGCs preferentially activate LC ^27^. Our findings also are consistent with a recent report that arousal-like behavior (linked by multiple previous studies to LC^2^) is increased in mice when ipRGCs are stimulated ^6^.

However, our findings also indicate a non-LC mechanism for the observed relative weight-gaining effects of PRAM stimulation—most likely driven by ipRGCs. These findings add to the recent, but expanding literature indicating an association between ipRGCs and metabolism. Mice null for melanopsin (Opn4-/-) mice show extreme weight loss following ketogenic diet challenge^28^. Constant darkness reduces normal circadian intestinal oscillations^29^, possibly by regulation of gut microbiota via ipRGCs^30^. Alternatively, ipRGCs may affect eating behavior via indirect input onto feeding-relevant regions of hypothalamus.

Our results indicate that PRAM stimulation also can prevent apoptotic and physiologic dysfunctions in LC-NA neurons. As loss of LC neurons is associated with several neurodegenerative disorders ^31–33^, our findings indicate a novel approach to better understand and prevent degenerative brain disease that underlies a variety of mental disorders. Our findings additionally indicate an important modulatory role of ipRGCs on LC function, indicating that PRAM activation can both prevent loss of LC neurons as well as improve neuropsychiatric dysfunction by activation of remaining LC neurons, compensating for a population that was previously lost. Recent results support this latter possibility for memory in an Alzheimer’s/Down’s model^34^.

Our findings point to a possible therapeutic route for the treatment of depression or other diseases associated with LC dysfunction, whereby the DREADD gene is delivered via intraocular injection, and daily DREADD-activating ligands are used to then activate the PRAM pathway. Activating LC via the PRAM pathway may have benefits over directly activating LC by local transduction of LC neurons by DREADDs.

We previously reported that CNO-mediated activation of hM3Dq-DREADD transduced into LC neurons induced bouts of immobility during a patch foraging task ^35^, similar to prior results with optogenetic LC activation^36^. Although the mechanism for such stimulation-induced immobility is unknown, it may reflect hyperactivation of LC neurons which induces anxiety or other maladaptive responses ^37^. PRAM activation is a more natural and nuanced means of increasing LC activity within the physiological range, somewhat akin to the activating effects of light on LC neurons ^38^. Thus, the minimally invasive method presented here is a promising approach for activating LC in both the laboratory and the clinic and may assist in allowing DREADD-based technology to evolve from an experimental to a clinical technique.

## SUPPLEMENTARY MATERIAL

### METHODS

#### Animals

Male Sprague-Dawley rats (250-300g) were single housed in transparent cages with access to food and water *ad libitum*. All animals were housed in a 12h: 12h reverse light / dark cycle (DL) prior to beginning the experiment. Four weeks following viral transfection of retinae (to allow time for DREADD expression), rats were housed in continuous darkness (DD) or DL for 8 or 10 weeks. Control and experimental animals were age-matched. All experiments were performed in compliance with Rutgers Animal Care and Use Committee guidelines, and procedures were approved by the Rutgers Institutional Animal Care and Use Committee.

#### Retinal transduction with hM3Dq vector

At ~8wks of age, rats were anesthetized, and pupils were dilated with 1% tropicamide ophthalmic solution (Akorn) and 1% cyclopentolate hydrochloride ophthalmic solution (Akorn). Either AAV2-hSyn-hM3Dq- mCherry (Experiment 1) or AAV-PACAP-hM3Dq-mCherry (Experiment 2) were bilaterally injected into the vitreous of both eyes (1µl each), using a 32 gauge needle (Hamilton Co., Reno NV) to target retinal ganglion cells (RGCs). The injection was made 1mm posterior to the temporal limbus, ~1.5mm deep, angled at ~45 °towards the optic nerve. Control animals were injected with AAV2-hSyn-GFP in an identical manner. CNO (2mg/kg, ip) dissolved in 3% DMSO was used to activate the hM3Dq receptor. Animals housed in DD were subjected to daily injections of CNO beginning from the first day in DD until the completion of behavioral tests. CNO was not present during these tests, so that our behavioral testing assessed the effect of preceding chronic, rather than acute hM3Dq PRAM activation.

#### Ablation of ipRGCs

We designed a ribosome inactivating protein (saporin, toxin) conjugated to a melanopsin antibody (Mel-SAP) to selectively ablate ipRGCs (Advanced Targeting Systems). Mel-SAP thus acts as a hybrid protein, combining the binding specificity of the melanopsin antibody with the cytocidal properties of saporins. At ~8wks of age, rats were anesthetized, and pupils were dilated as above. Either Mel-SAP or PBS was bilaterally injected into the vitreous of both eyes in an identical manner to that described above for the viral transduction of hM3Dq into retinae. After waiting 2wk for ablation, animals were housed in DL for 8wks until the completion of behavioral tests.

#### Quantification of melanopsin+ cell ablation

Whole retinae were gently separated from vitreous and retinal pigment epithelium. Tissue was incubated in 2% NDS to reduce non-specific antibody binding. Primary antibody was directed against melanopsin (rabbit *α* melanopsin: 1:500; PA1-781; Thermo Scientific, Waltham, MA) and combined with 2% NDS for incubation overnight. The next day, retinal tissue was incubated in Alexa Fluor 594-conjugated donkey anti-rabbit for 2 hrs. Retinae were rinsed in PB, flat-mounted onto glass slides and coverslipped using Citifluor AF1 Mountant Solution (EMS, Hatfield, PA). Images of each entire flat-mounted retina were obtained (ZenPro software). Offline high-definition images were used to count labeled cells using ImageJ software. Cells were counted and simultaneously marked such that each cell could only be counted once.

#### Recordings of Locus Coeruleus

Locus coeruleus (LC) activity was recorded using either a 25um diameter wire or a pipette while the animal was under 2% isoflurane anesthesia. Rats were dark-adapted for at least 12h before recording, and eyes were masked with blackout rubberized fabric (BK5; Thor Labs) throughout the surgery and recording procedure. The initial penetration was made based on anatomical landmarks and confirmed by known physiological characteristics of LC activity to a brief foot shock: phasic discharge of 2 or 3 action potentials (10–20 Hz), followed by sustained suppression of spontaneous activity (200–500 ms). A single unit was isolated and its activity was recorded for ~10 minutes. Following recording each single unit, the electrode was advanced or relocated to find another LC unit to record. After 1-2 hours of recording, consisting of 1- 3 penetrations, 2mg/kg CNO was injected ip. After 30 minutes, recording from LC units was repeated, following the same procedure described above. Cells were sampled both before and after CNO administration when possible, but populations of cells recorded before vs after CNO also were compared.

### Behavioral Tests

#### Saccharin Preference Test

Rats were habituated to two bottles placed in the home cage over two consecutive days. Access to both bottles containing either water or 0.1% saccharin dissolved in water was given through sipper spouts. The left-right position of the bottles was alternated every 12h. The following day, bottles containing water and 0.1% saccharin were reintroduced to the home cage, this time for 2 h. The left-right position of the bottles was alternated after 1h. Preference for saccharine were calculated as a percentage of the total liquid intake using the formula: saccharin intake/(water intake+ saccharin intake) x100.

#### Forced swim

Animals were individually placed in a cylindrical Plexiglas tank (38.5cm high × 30.5cm diameter) filled with tepid water (25-27 °C). On the initial test day, animals were subjected to the FST for 15min. Twenty-four h later, rats were subjected to an identical test, but for 5min. A video camera mounted at the front of the cylinder recorded activity. An observer, blind to experimental conditions, scored the animals’ behaviors on the second test according to three criteria: swimming, climbing and immobility. Immobility was defined as the rat floating in the water without struggling, and only making movements necessary to keep its head above water. For simplicity, only immobility data is presented. To avoid confounding other behavioral tests by stress induced by the FST, the FST was performed following the completion of all other behavioral tests.

#### Elevated Plus Maze

The elevated plus maze (EPM: Med Associates, St Albans, VT) consisted of four arms measuring 50 cm x 10 cm, and was elevated 50cm above the ground. Testing occurred at noon, approximately 20h following the last CNO injection in groups that received daily CNO. Rats were individually placed on the center square of the EPM. The time spent in each arm was automatically recorded by a computer monitoring photobeam crosses for 15 minutes. After each trial, the EPM was cleaned and dried.

#### Locomotor Testing

Rats were placed in locomotor chambers (42cm x 42cm x 30cm) fitted with SuperFlex monitors (Omintech Electronics Inc, Columbus, OH), a 16 x 16 infrared light beam array for the x/y axis, and a 16 x 16 array for the z axis. Activity was automatically recorded over a 2hr period by Fusion SuperFlex software.

### Quantification of Apoptosis in Locus Coeruleus

PARP immunoreactivity was assessed by incubating 40 μm-thick sections in rabbit antiserum against the p85 fragment of PARP (1:100; Promega) overnight. To localize LC, caudal brain sections were concurrently incubated in mouse anti-TH (1:1000; Immunostar). Secondary antibodies were a cocktail of Alexa Fluor 594-conjugated donkey anti-rabbit (1:500; Jackson Laboratories) and Alexa Fluor 488 donkey anti-mouse (1:500; Jackson Laboratories) in which tissue was incubated for 2h.

Images of LC were obtained using a Zeiss Axiozoom microscope attached to a monochromatic camera (ZenPro software). Offline high-definition images were used to count labeled cells using ImageJ software (NIH). Three bregma levels were (rostral, mid, caudal; −9.4 mm, −9.7 mm, and −10.0 mm from bregma) were averaged to determine the overall apoptosis index of each animal

### Quantification of DBH+ boutons

Prefrontal cortex (PFC) sections (40μm-thick) were incubated in primary antibody directed against DβH (mouse anti-DβH 1:1000; Millipore; MAB 308) for 48 h at 4° C. The following day tissue was incubated in donkey anti-mouse (1:500); Jackson), amplified with avidin biotin complex (ABC 1:500; Vector Labs) and visualized with DAB Slices were mounted onto glass slides, washed and dehydrated through alcohol and xylenes, and coverslipped with DPX (Sigma).

Stained boutons were quantified to assess noradrenergic (NA) innervation of PFC. The number of DβH+ boutons in PFC of each brain was estimated using unbiased stereological methods (Stereologer; SRC, Chester, MD). The dissector technique was used to randomly count boutons in a defined area (frame size: 2000 µm2, frame height: 15 µm, guard height: 2 µm, frame spacing: 100 µm) in the cingulate-prelimbic cortex (3.70 mm rostral to bregma). The total number of DβH+ boutons was calculated by the software as: n = number of boutons counted × 1/section sampling fraction × 1/area sampling fraction × 1/thickness sampling fraction. All analyses were performed blinded to experimental group.

### Data analysis

Prism 9.0 (GraphPad Software) was used for all statistical analyses. Values were compared using one-way or two-way ANOVAs with Holm-Sidak post-hoc tests as appropriate. In cases where data were not normally distributed, appropriate non-parametric analyses were used. A P value of <0.05 was considered statistically significant. Error bars represent means ± s.e.m. All *n* values represent individual rats.

## SUPPLEMENTARY FIGURES

**Supplementary Figure 1.**
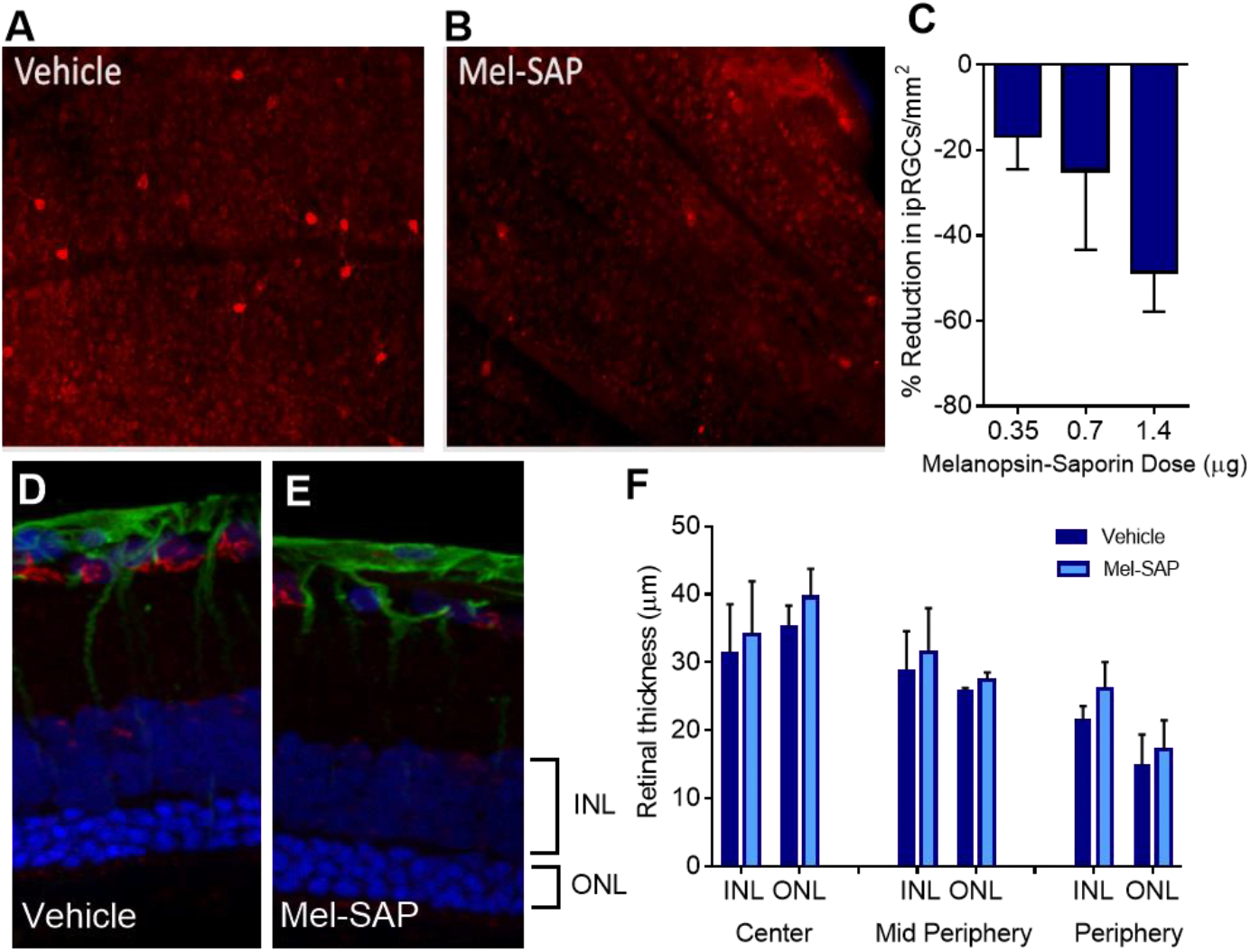
*Melanopsin-Saporin intravitreal injection leads to selective reduction of melanopsin containing retinal ganglion cell*s. (A) Melanopsin staining of retinal ganglion cells (RGCs: flatmounted) in animals that received vehicle. (B) Melanopsin staining of animals that received Mel-SAP. (C) The highest dose of MEL-SAP (1.4 ug) led to ~50% reduction in melanopsin RGCs. This dose was subsequently used for behavioral testing. (D) Retinal cross sections of animals that received vehicle and (E) Mel-SAP show that the knockdown was specific to melanopsin containing RGCs (shown in red). (F) Thickness of the inner nuclear layer (INL) and outer nuclear layer (ONL) was measured in the center, mid periphery and periphery. No differences were seen between groups, indicating robust survival of all remaining non-melanopsin cells of the retina.

